# RANDOMIZE: A Web Server for Data Randomization

**DOI:** 10.1101/2020.04.02.013656

**Authors:** Agaz H Wani, D Armstrong, Jan Dahrendorff, Monica Uddin

## Abstract

**Summary:** DNA methylation microarray data may suffer from batch effects due to improper handling of the samples during the plating process. RANDOMIZE is a web-based application designed to perform randomization of relevant metadata to evenly distribute samples across the factors typically responsible for batch effects in DNA methylation microarrays, such as row, chips and plates. Randomization helps to reduce the likelihood of bias and impact of difference among groups.

**Availability:** The tool is freely available online at https://coph-usf.shinyapps.io/RANDOMIZE/ and can be accessed using any web browser. Sample data and tutorial is also available with the tool.

## 1. Introduction

DNA methylation is a critical type of epigenetic modification that typically occurs in CpG-rich regions in the mammalian genome and is associated with regulating gene expression (Sharma et al., 2010; Moore et al., 2013). Previous studies have revealed a strong association of change in DNA methylation with various diseases such as cancer (Karpiński et al., 2008; Feinberg and Irizarry, 2010) and post-traumatic stress disorder (PTSD) (Uddin et al., 2018).

High throughput microarray technology has made it possible to measure methylation levels of thousands of probes simultaneously in an inexpensive manner. The microarray-based Illumina Infinium MethylationEpic Bead (Epic 850k) Chip has become a useful and standard tool for epigenome-wide DNA methylation profiling. The technology interrogates over 850,000 selected methylation sites (CpGs) per sample at single-nucleotide resolution, including >90 % of the CpGs from the Illumina HumanMethylation450 Bead Chip and an additional 413,743 CpGs (Ruth Pidsley et al. 2016). Each Epic 850k chip can accommodate eight samples, and each 96 well plate has 12 chips for 96 samples in total. Thus the samples in large studies are often assayed across different chips and plates and processed in different batches. Accordingly, there could be a lot of non-biological variations due to experimental factors such as conditions in the laboratory, time of the experiment, reagent differences, personnel differences in preparing the samples, and chip position(row). This variation may give rise to batch effects (Yan et al., 2012; Harper et al., 2013) that affect the methylation level of different probes. Batch effects are a significant issue and can lead to spurious and inaccurate results and reduction in power to detect real biological differences (Akey et al., 2007).

Batch effects are difficult to remove entirely during the normalization process following data collection. Even the effectiveness of the advance techniques like ComBat (Johnson et al., 2007) to adjust for batch effects depends on the study design. Harper et al. (2013) found that even powerful techniques such as ComBat could not wholly remove batch effects when the samples are not randomized across chips, thus leading to false detection of differentially methylated probes. Since batch effects can’t be disposed entirely of from even a perfectly designed study, Hu et al. (2005) emphasized that careful study design is crucial to address batch effects and other technical artifacts. For example, in a case-control study, the cases and controls should be uniformly distributed across the factors considered to be responsible for a batch effect. This can help to avoid problems such as those identified by Liu et al. (2013), who found a surprising relationship between methylation data and assay date due to the unbalanced distribution of cases and controls on those dates.

All this tells us that it is essential to randomize the samples to reduce the likelihood of bias. Random assignment of samples to row, chip, and plate ensures that each sample has the same probability of being attached to a particular chip and thus satisfies the requirement of uniform distribution of the data. Randomizing the samples can be a tedious, error-prone, and time-consuming task when dealing with hundreds of samples. According to our knowledge, there is no tool existing to perform randomization. To facilitate this process, here we present a web-based tool that helps users to randomize samples in a user-friendly and efficient way. The tool can randomize hundreds of samples within a matter of a few seconds and is available online and free to use.

## 2. Usage

We developed the tool RANDOMIZE with the primary purpose of providing a user-friendly, GUI-based tool for biologists to perform randomization of the metadata. The tool is very simple to use. As of now, it is compatible with randomizing samples on 96 well plates. There is an option to balance randomization on various factors such as case-control, male-female, etc. The user needs to feed a CSV file as an input where the data is available across different columns. The user selects the columns on which the randomization needs to be balanced and then submits the job for processing. There is also an option to constrain known controls (or duplicate samples) at any location on the chips.

Once the sample assignment has been randomized, the user can perform an exploratory analysis of the randomization results. Many plots are available for exploratory analysis and to check the goodness of randomization, including Sunflower, Violin and Density plots. The final study design is available to download for further usage.

## 3. Analysis

For the assessment of functions and illustration of tool utility, we have used sample data with 750 samples. The sample data is available with the tool at the above given link.

Go to the “Analysis” tab to start randomization. On the right side is the “Randomization**”** panel, shown below in Figure 1, where you can browse your computer in order to locate your metadata file and upload the metadata file in a CSV file format. Successful uploads will be indicated as “Upload complete”. It is important that your metadata file include columns labelled as “ParticipantID**”** and “SampleID**”**.

**Figure 1:**
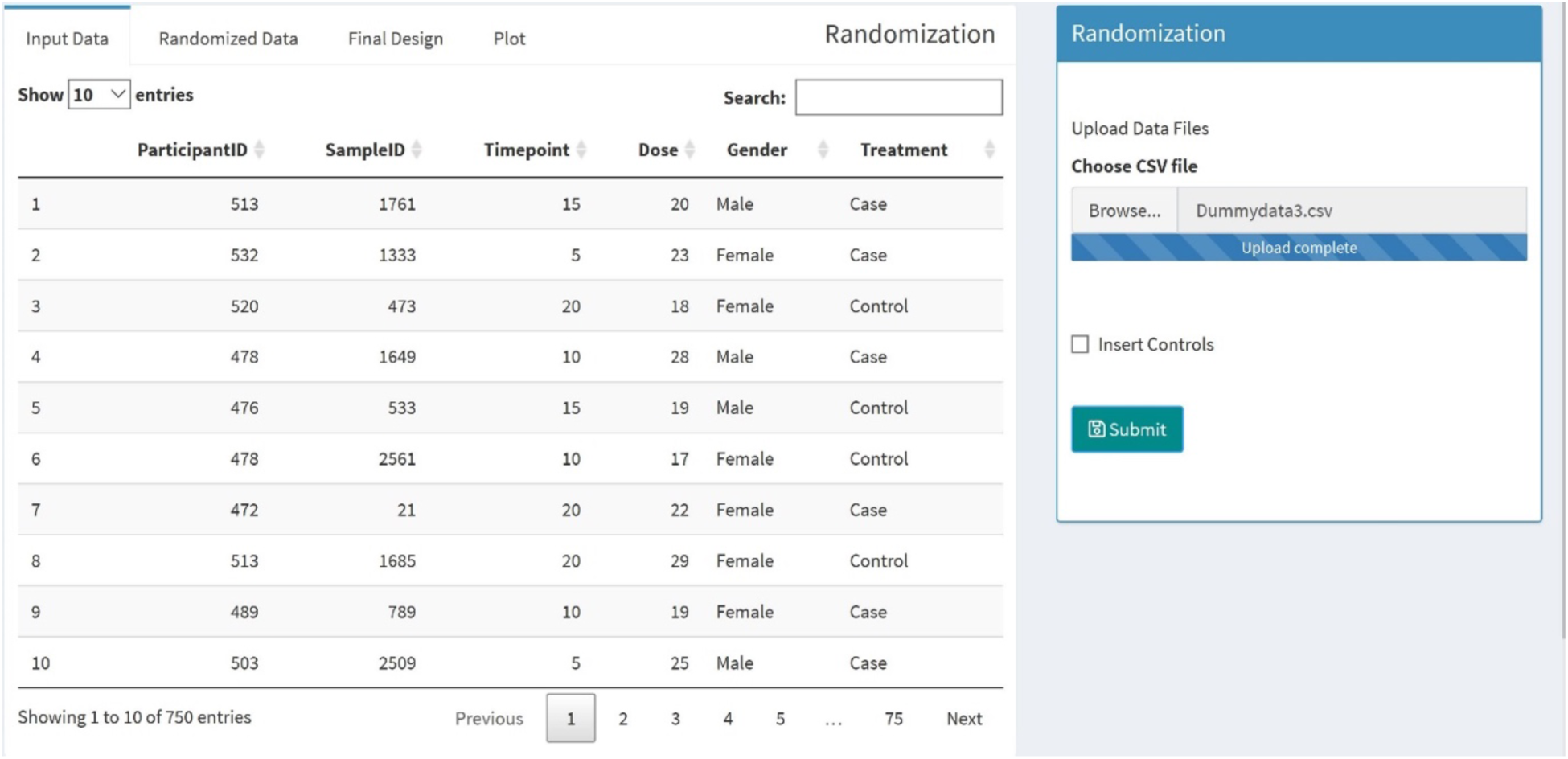
shows input data.

Your data should show up in the “Input Data**”** tab on the top left. In the Randomization panel, you can also check the box “Insert controls” to manually add known controls to your analysis as shown in Figure 2. Controls can then be added on individual chips. No controls are inserted by default.

**Figure 2:**
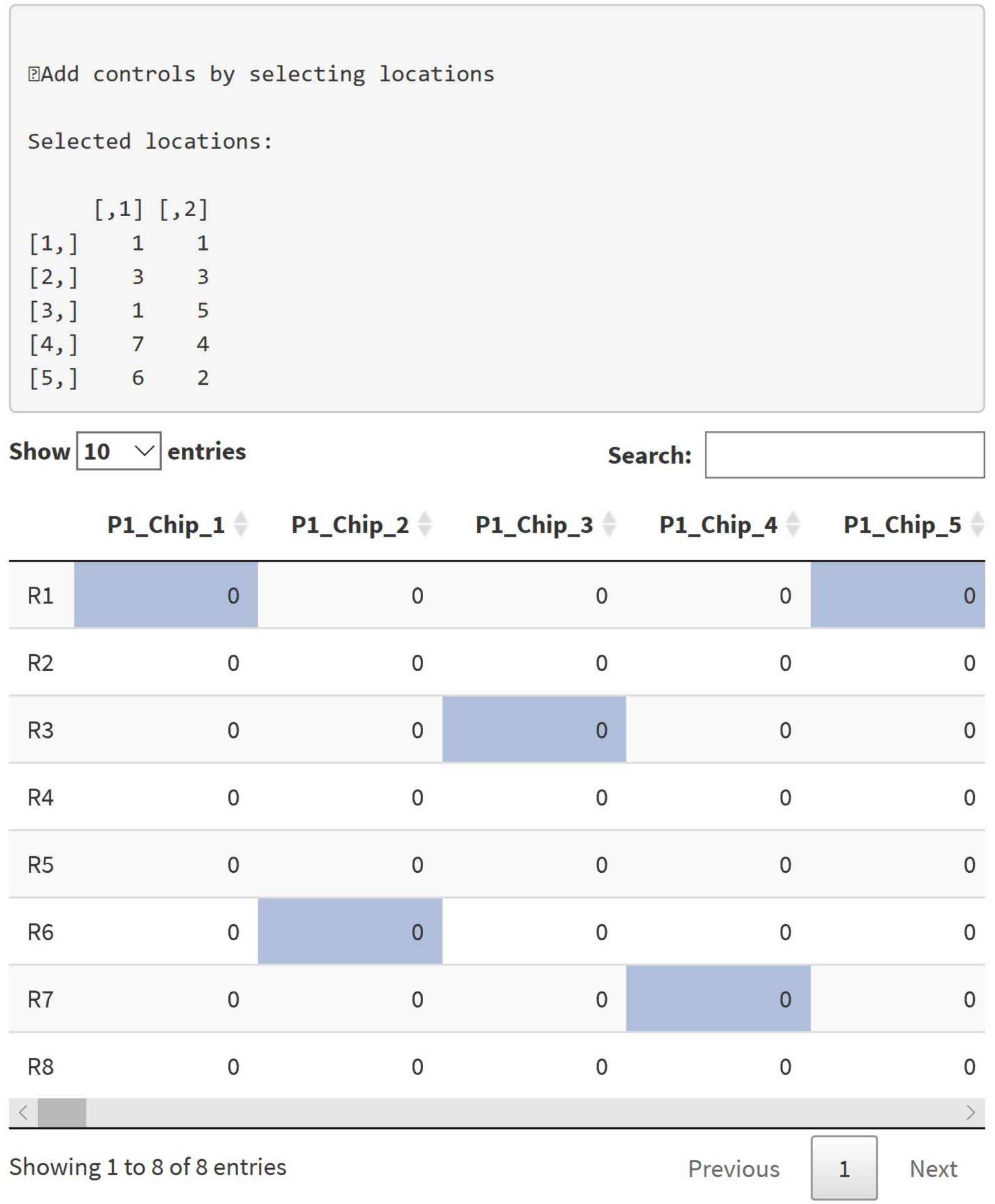
shows places for controls.

By clicking on the columns, you are able to select the columns you would like to randomize (shown below in Figure 3). If you hit “Submit” on the randomization panel without selecting any columns, you will receive an error message: “Please select columns for randomization by clicking on desired column(s)”.

**Figure 3:**
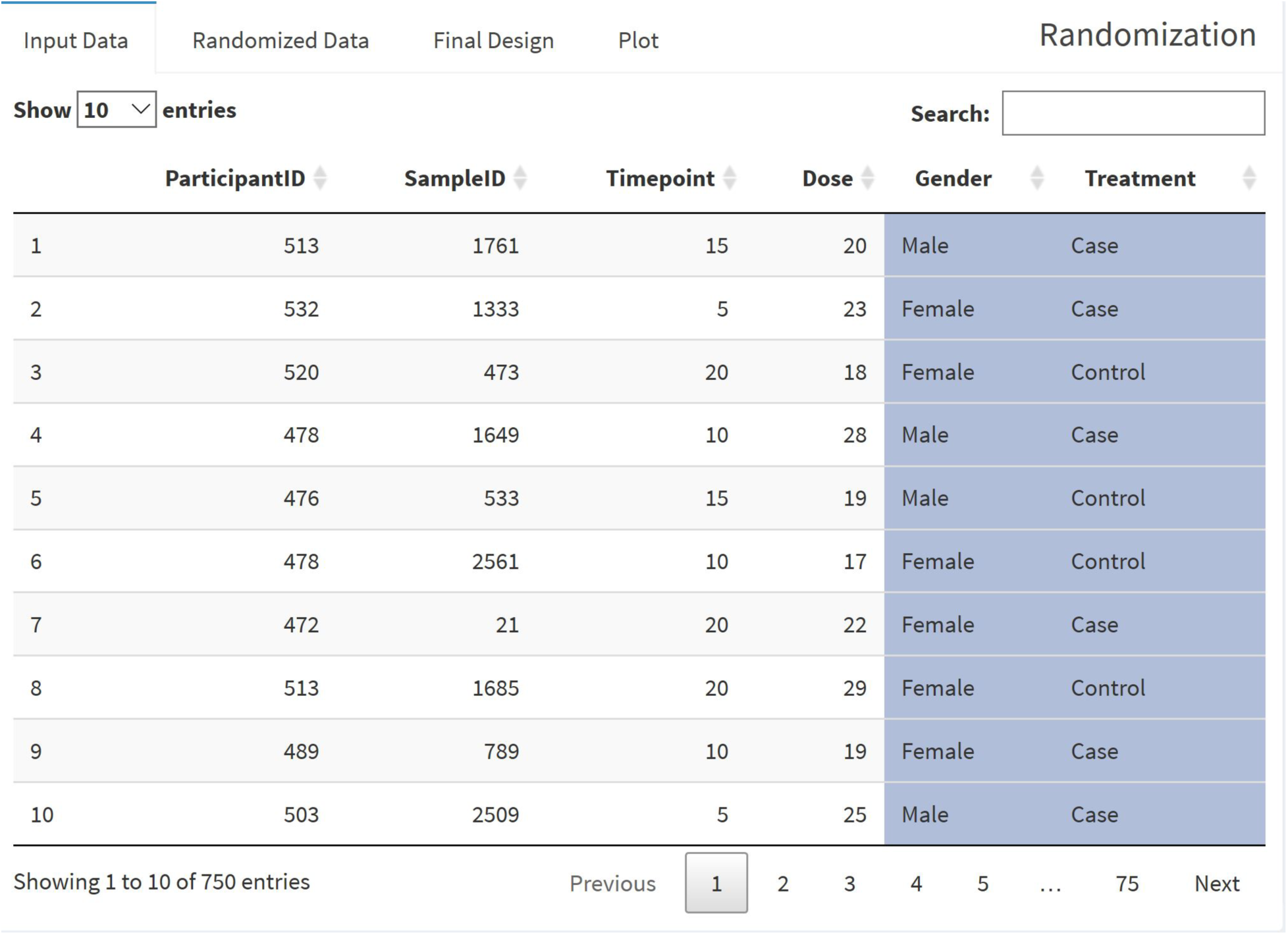
Selected columns of interest for randomization are highlighted in blue.

After selecting the columns of interest for randomization click on the “Submit Button” located on the bottom of the Randomization Panel to submit the job for processing. Once the job is processed, in the “Randomized Data” tab (next to the “Input Data**”** tab) you can take a look at your randomized data of your selected items, as shown below. Your previously selected controls are excluded from the randomization and still in the location you have selected beforehand. The controls are shown as zeros, as in Figure 4.

**Figure 4:**
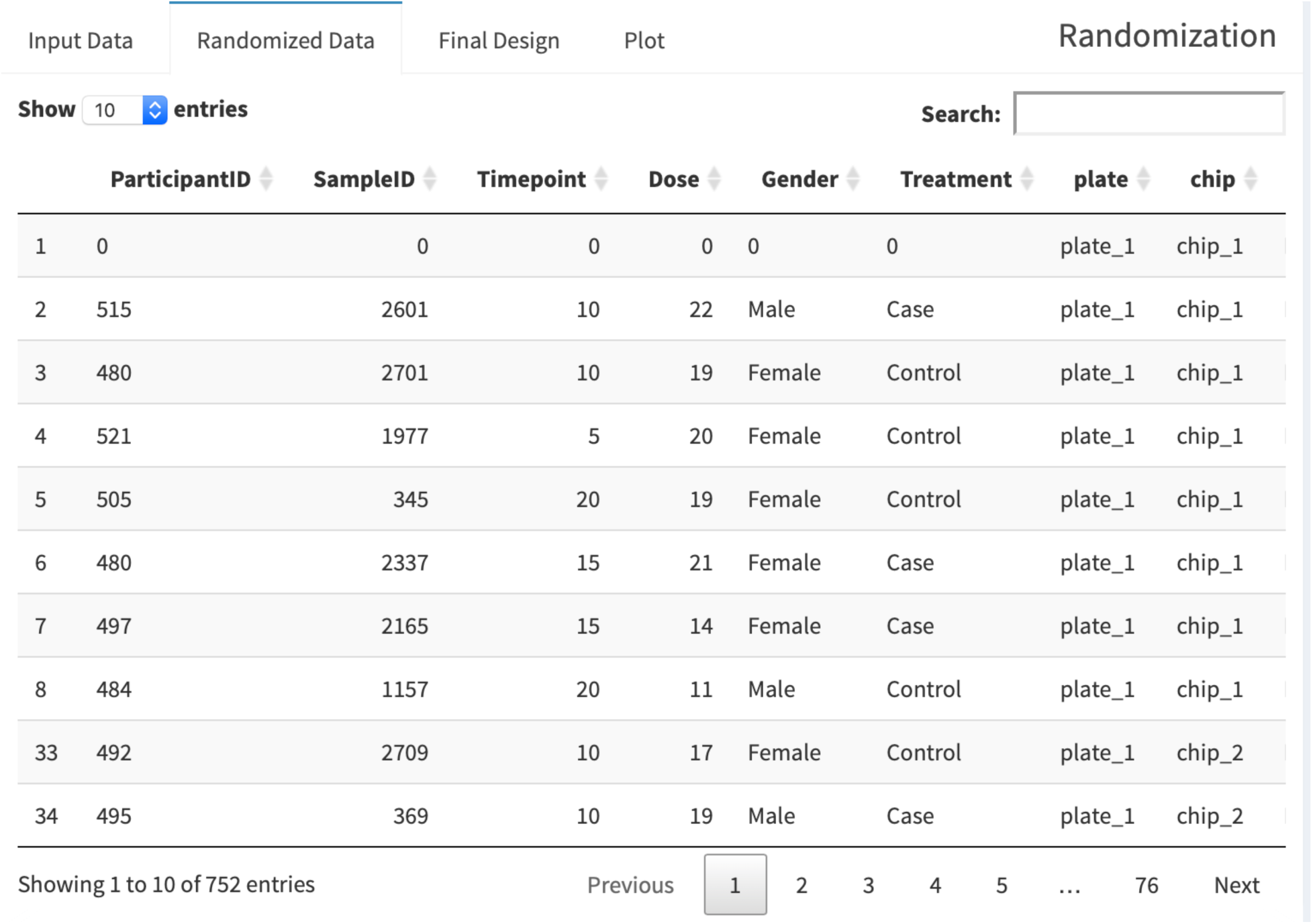
shows randomized data.

The “Final Design**”** tab adjacent to the “Randomized data**”** tab shows you the final design of your randomized data. The “Display Final Data**”** tab lets you view your final design, one plate at a time. The design of the first plate is available to view by default. The “Plot” tab eventually shows you your plotted data. You should select the columns of interest by clicking on them before you move on to the “Plot” tab. In the “Plots**”** selection on the left you are able to choose between various plots. The “Plot labels**”** option lets you select a title for your plot and label the x- and y-axis.

As an example we illustrate the goodness of randomization using the sample dataset and sunflower plot. The sunflower plot is used to display bivariate distribution. Each petal on the sunflower plot represents an observation(sample). The “ParticipantID**”** column in our sample dataset denotes the participant ids; each participant has one or more samples in the range of 1-23. There are 112 unique participants in the dataset. For 750 samples, eight samples on one chip, we need 94 chips in total. An ideal randomization would be that no two or more samples from the same participant are on the same chip; however, the number of chips is less than the number of participants, so it is evident that some samples from the same participant will be on the same chip. The black dots in Figure 5 denote unique samples. If two samples from the same participant are on the same chip, a petal, as shown in red, is added on the black dot. For two duplicates, two petals are added, and so on. The plot indicates proper randomization of the data — for example, for the participant which has 23 samples, all the samples are sent to different chips. Only some chips have two samples from the same participant id. Similarly, the randomization of participant ids on plates is shown in Figure 6.

**Figure 5:**
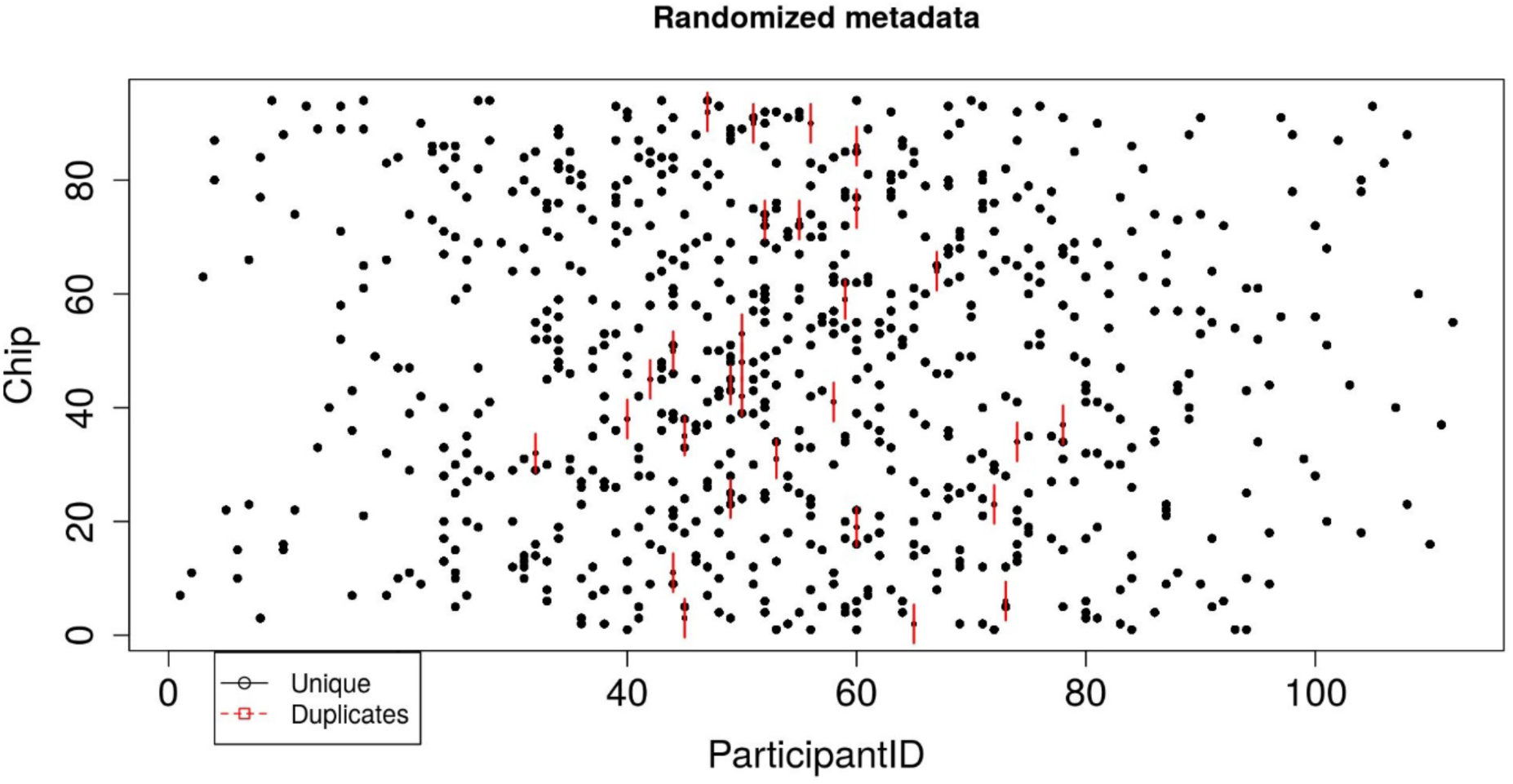
Sunflower plot showing the metadata distribution (here, participantID) across Chips.

**Figure 6:**
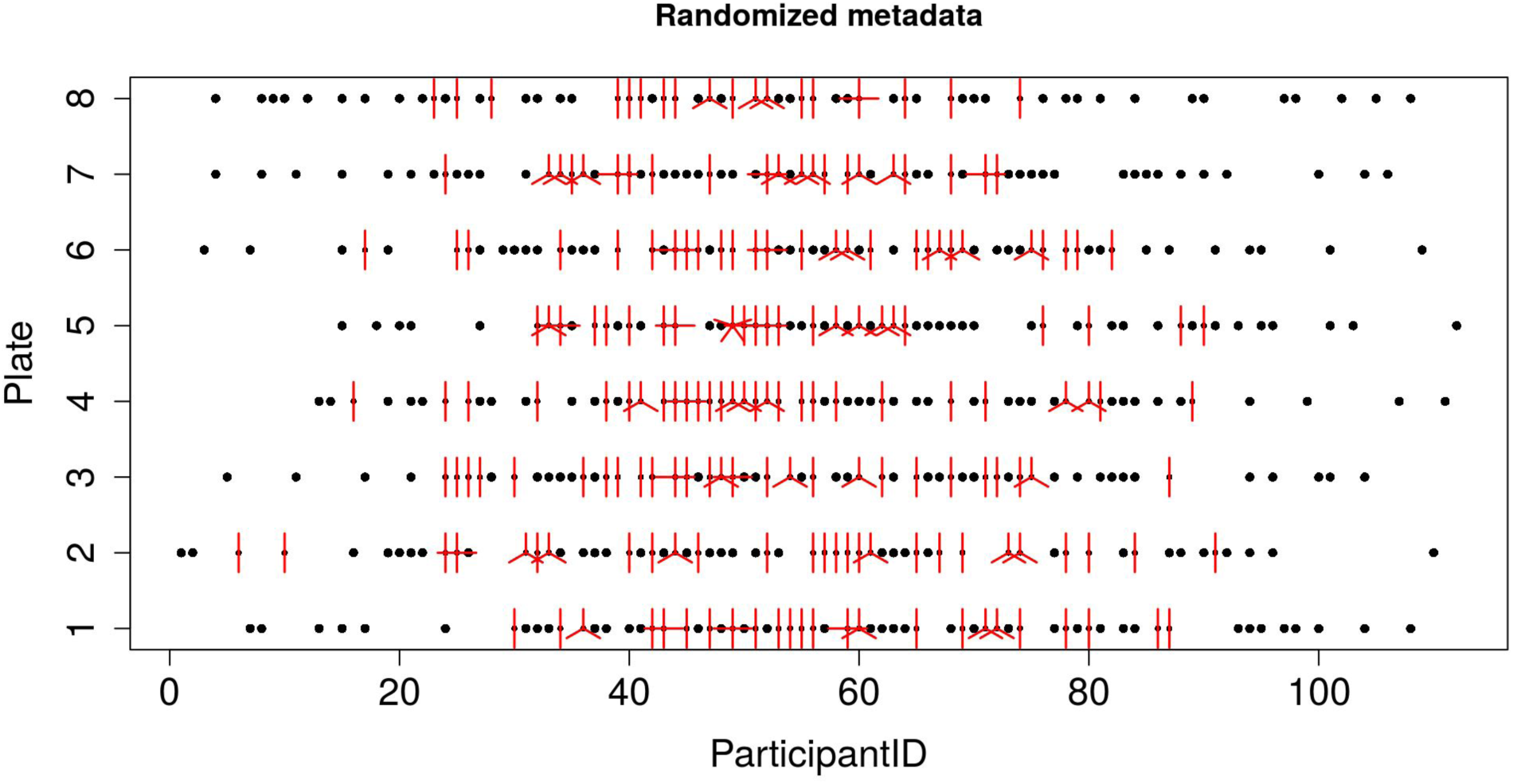
Sunflower plot showing the metadata distribution (here, participantID) across Plates.

The “Download” tab lets you download your randomized data. In the “Randomized Data**”** tab, you are able to download your “Randomized Data**”** and “Final Design**”** in a CSV file format while an individually downloaded “Plot**”** will be in a PNG file format.

## 4. Implementation

The graphical user interface of the tool was designed and implemented using the R library (shiny 2013), and the methodology was applied using R 3.6.1 (R 2017) and RStudio 1.0.44. The tool can be run on any operating system, including Windows, Linux, and is available using any web browser (best viewed on Firefox, Google Chrome, and Safari).

## 5. Conclusion

High-throughput DNA methylation arrays are susceptible to bias facilitated by batch effects and other technical noise that can alter DNA methylation level estimates. RANDOMIZE is a user-friendly web application that provides an interactive and flexible graphical user interface (GUI) to randomize relevant metadata. Using this tool will minimize chip and position mediated batch effects in microarray studies for an increased validity in inferences from methylation data. The tool is very helpful for biologist to perform randomization of test samples and insert controls in the data.

## Acknowledgements

This work has been supported by National Institutes of Health, grants 1R01MD011728 and R01MD011728-S2

